# The First Report on Cryptococcus Profiles of Isolates from Patients Attending Dr George Mukhari Academic Hospital, South Africa

**DOI:** 10.1101/620930

**Authors:** Elliot Zwelibanzi Jiyane, Mis Leah Nemarude, Maphoshane Nchabeleng

**Affiliations:** Sefako Makgatho Health Science University, Oral Health Center, School of Dentistry, Department of Oral Microbiology, Pretoria, South Africa email address; Sefako Makgatho Health Science University, Microbiological Pathology, School of Medicine, Pretoria, South Africa email address; Sefako Makgatho Health Science, University Microbiological Pathology, School of Medicine, Pretoria, South Africa email address

**Author notes:** Corresponding author: Jiyane Mailing address: Medunsa Campus, PO Box CP30, Sefako Makgatho Health Science University, 0204.

**Keywords:** cryptococcosis, serotypes

## Abstract

**Introduction:** Cryptococcosis is a fungal opportunistic infection that is vastly diagnosed among immune-compromised patients. Reduced susceptibility on commonly used antifungals is of concern. In the communities served by Dr. George Mukhari Tertiary (DGMT-Laboratory) Laboratory is not available.

**Methodology:** E-test method was used to determine if isolates with reduced susceptibility to antifungals fluconazole, voriconazole and amphotericin-B had emerged. A multiplex Polymerase Chain Reaction (PCR) method was used to further identify serotypes that are circulating at Dr. George Mukhari Tertiary (DGMT-Hospital) Hospital.

**Results:** E-test strips were interpreted as resistance, intermediate or susceptible in relation to each serotype identified. Of the 50 incident isolates tested, 100% were inhibited by both voriconazole and amphotericin-B. Fluconazole was resistance to 50% of incident isolates.

**Conclusion:** *C. neoformans* serotype A is the predominant serotype in the area served by DGMT-Laboratory, accounting for 96% of the isolates. It is important for public health to continuously monitor resistance emergence.

## BACKGROUND

Cryptococcosis is amongst the leading and life-threatening opportunistic infection ^[1]^. The disease is caused by *Cryptococcus neoformans (C. neoformans)* and *Cryptococcus gattii (C. gattii)*, which are vastly diagnosed among immune-compromised patients^[2]^. Mainly *C. neoformans*, with three recognized serotypes which are acknowledged as serotype A, serotype D and the hybrid-AD^[3]^. Previously, these serotypes were identified and differentiated by the phenotypic approach^[4]^. Lately, are identified by PCR assays^[5]^. More methods of molecular assays are used to classify *C. neoformans* serotypes^[6]^. The classification is based on antigenic metamorphoses in the polysaccharide capsule associated with virulence factors ^[3,7]^. Prevalently found circulating globally is serotype A, whereas serotype D and AD hybrid circulating in truncated numbers^[1–3]^. Serotype A account for more than 90% cases of cryptococcosis in South Africa owing to the extraordinary occurrence of HIV/AIDS^[2,8,9,10]^.

To treat cryptococcosis, the most widely used antifungal agents are amphotericin-B, flucytosine, voriconazole and fluconazole^[9,11]^. Amphotericin-B is used as the first line treatment but limited by toxicity that requires laboratory monitoring, voriconazole is limited to private sectors in Africa^[9-12]^. The amalgamations of antifungals are recommended for induction but flucytosine is not available in resource-poor countries^[12]^. Unfortunately, these are countries with a high incidence of cryptococcosis^[10-12]^. All these antifungals are also limited by emerging resistance mechanisms such as the antigenic polysaccharide capsule tolerance, the mating gene types, the acid tolerant abilities, and spores switching^[12]^. Globally, 20-58% of resistance cases are reported on cryptococcosis by means of diverse studies, focusing on fluconazole^[8-10,13,14]^. Emerging resistance on the other cryptococcosis antifungals was not reported^[9,15]^. Furthermore, intrinsic and acquired resistance mechanisms are all associated with Cryptococci and the drugs proneness to those resistance mechanisms ^[12,16-18]^

The widespread use of fluconazole may lead to the emergence of reduced susceptibility^[19,20]^. Thus the development of resistance to fluconazole is devastating to the treatment of cryptococcosis, and it is necessary to know if there is cross-resistance with voriconazole which could be an alternative agent. It is important for institutions to monitor for changes in susceptibility profiles of isolates circulating in their areas in order to update the treatment regimens. Our aim in this study was to identify circulating serotypes and determination of the susceptibility profiles of Cryptococcus isolates to fluconazole, voriconazole and amphotericin-B antifungals form clinical specimens sent to the DGMT-Hospital NHLS.

## METHODS

### Ethical consideration

Ethics were sorted from Medunsa Research Ethics Committee. Permission to obtain isolates was sorted from the DGMTL and NHLS managers. Clinical isolates were delinked from identifiers to ensure confidentiality.

### Sample size

Epi Info version 3.5.3 (Centre for Disease Control and Prevention) was used to calculate the sample size. The required sample size in this study was 50. This was calculated at an estimated frequency of 50%, power of 80% and the confidence interval of 95%.

### Demographics

Demographic data including age, sex and clinical diagnosis of patients from whom the isolates were isolated was obtained from the Laboratory Information Systems (LIS) in the laboratory.

### Collection and storage of isolates

Clinical isolates were conveniently collected from the laboratory after processing for patient management was completed. The isolates were collected from February-July 2014, on a day to day basis until the sample size was reached. These isolates were already identified by the NHLS as ***Cryptococcus*** and stored in a - 4°C fridge.

### Sub-culture of isolates

The stored isolates were sub-cultured on Sabouraud dextrose agar (SDA) plate as described by Govender et al. 2011^[9]^.

### Confirmation and identification of Cryptococcus

Gram staining was done to confirm the morphology of yeast cell according to Chayakulkeeree (2007) description^[21]^. India ink (negative stain) was done as described by Ogundeji (2013) to verify the presence of capsule^[22]^. The isolates were further inoculated to urease broth media test in a slant position as a confirmation test according to Gazzoni (2014) methods^[23]^.

### Susceptibility Testing

After sub-culturing on SDA (**Figure 1**), susceptibility testing was achieved according to Clinical and Laboratory Standards Institution (CLSI) outlines of 2007^[24]^ and as described by Govender et al. 2011, 2013^[9,25]^.

**Figure 1:**
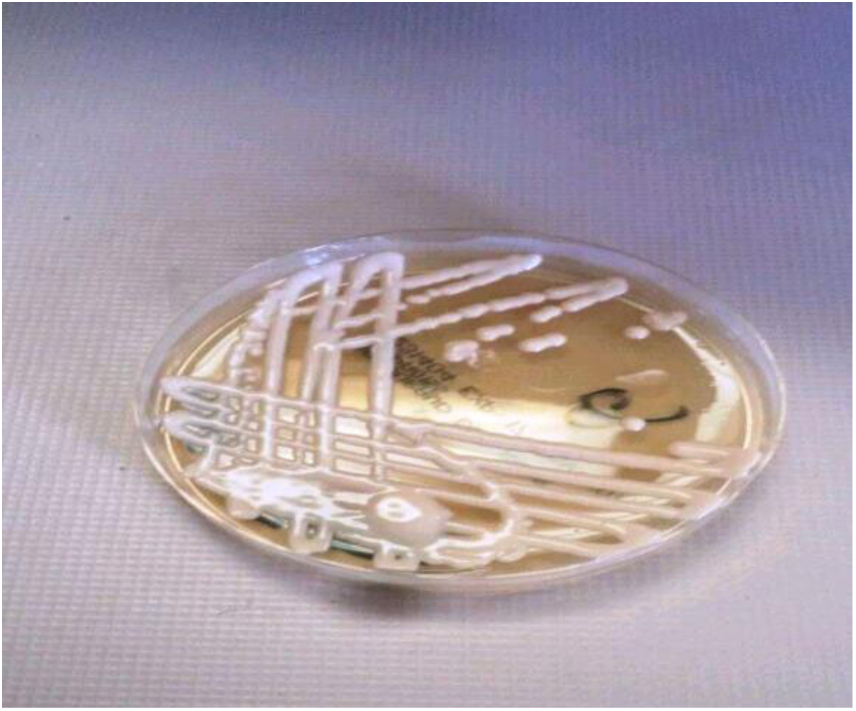
Mucoid colonies on SDA medium

### DNA Extraction and Sample Preparation for Multiplex PCR

Genomic DNA was extracted from the clinical isolates using the commercial kit (ZR fungal/bacterial DNA MiniPrep kit) in accordance with the manufacturer’s instructions (Zymo research group). The kit has been optimized for removal of PCR inhibitors and maximal recovery of pure DNA without RNA contamination. The extraction of DNA from the isolates was done using the protocol, “Biological liquids and cell suspensions”^[26]^.

### Primers selection

The primers used were synthesized by Inqaba Biotechnical Industries (Pty) Ltd, Muckleneuk, and Pretoria. The serotypes specific primers were designed to target the Mating - a gene and Mating - a gene of both serotypes A and D^[27]^. Primers targeting for genes confirming *C. neoformans* serotypes are listed in **table 3**.

**Table 1:**
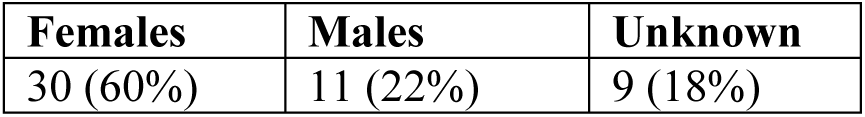
Demographics of the patients. Only 41 of the 50 patients from where the clinical specimens were sent had complete information from the laboratory information system.

**Table 2:**
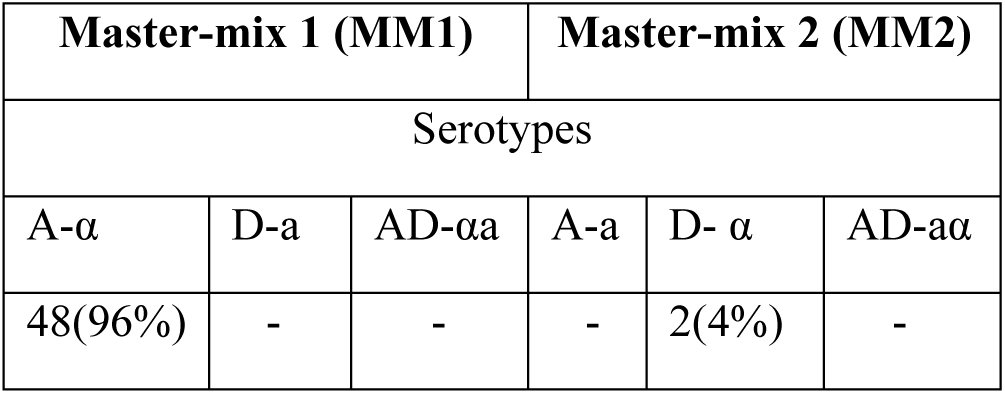
Molecular confirmation of serotypes. PCR for serotyping of *C. neoformans* was performed on all 50 isolates.

**Table 3:**
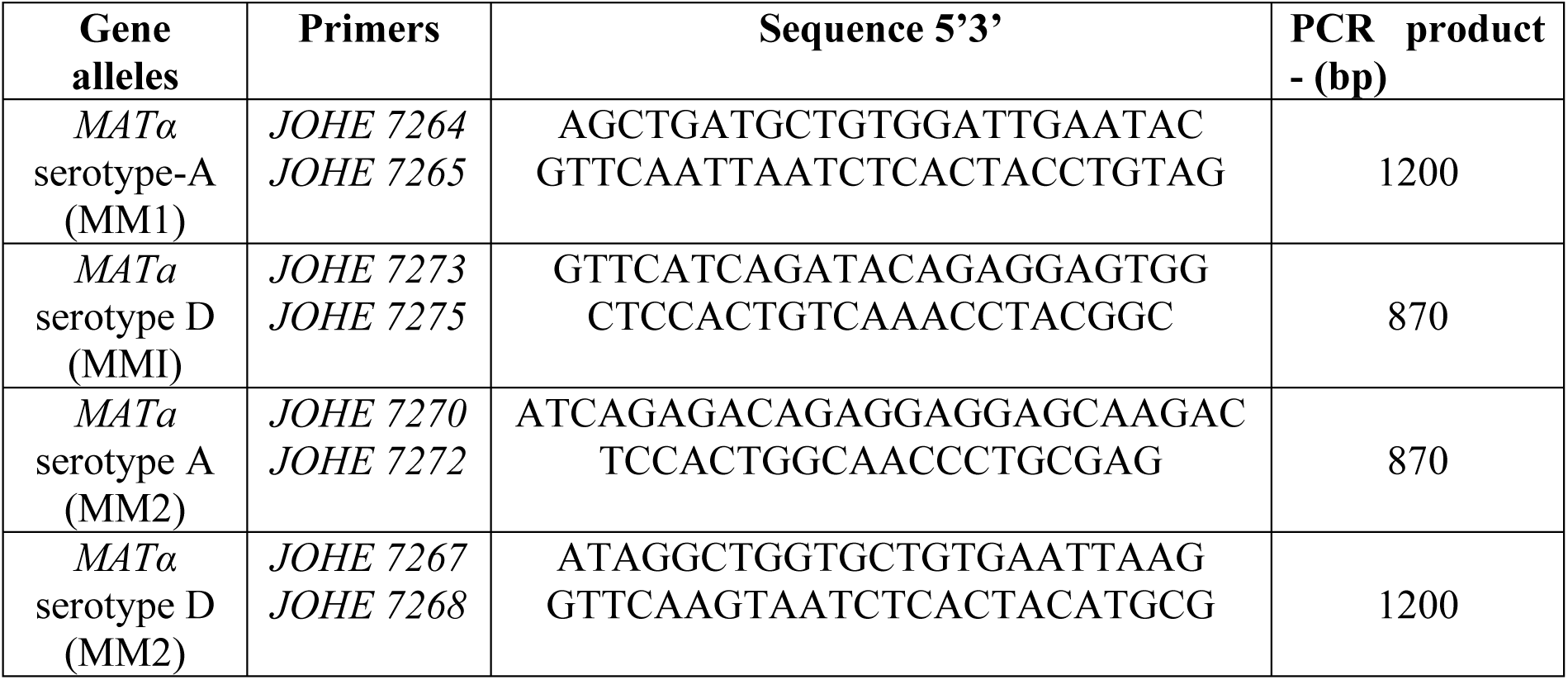
Combination and sequences of the primers used for the determination of serotype and mating type of *C. neoformans* by PCR multiplex alpha-Aa-D and a-Aalpha-D (N= 50)

### Amplification of genes

This was done on the extracted DNA using specific primers **(Table 3)**. Two master-mixtures (MM) were prepared. Reagents were obtained from Bioline Meridian Life Science Company (UK), each PCR assay was set-up with nuclease-free water as the negative control (Bioline, UK), and positive controls were not included due to financial constraints. MyTaq™ HS DNA-Polymerase (Bioline, UK) was used in the PCR reactions.

For each sample, a 50 μl reaction MM was prepared following the manufacturer’s instructions (Bioline, UK).

Briefly: 10 μL x MyTaq™ HS buffer, 1 μL of each of 2 primers, 5 μL of the template, 0.5 μL MyTaq™ HS DNA-Polymerase (Bioline, UK) (5 UμL), and 32,5 μL nuclease-free water. Two sets of MM were used, in MM1 contained alpha-A***a***-D primer set and the MM2 contained *a-Aalpha-D.* The 0.2 mL PCR tubes each containing 50 μL placed into a reaction was allowed to take place in a GeneAmp PCR System 9700 (MTHE 01326 PE Applied Biosystems) thermocycler for 3 hours; succeeding PCR temperatures as described by Saiki (1999)^[28]^.

### Detection of amplified products

Electrophoresis was performed on all samples using 2,0% agarose gel (Crystal TBE, Bioline, UK) for 40 minutes at 100 V, with ethidium bromide and UV transilluminator (Gel Doc™ EZ System). The 1 kb molecular weight marker (HyperLadder IV, Bioline, UK) was used in together with the amplified products. The photographic copy was taken using a Gel Doc EZ imager and the results were recorded as representative of serotype-Aα, Dα or A-a, D-a genes. For expected bands see **table 3.**

### Capturing of data

Microsoft Excel (Microsoft Office, 2010) was used to analyze data and the captured data was double-checked to ensure reliability; Epi Info version 3.5.3 (Centers for Disease Control & Prevention). Descriptive statistical analysis was performed based on ANOVA excel, 2010. Measures of central tendency and dispersion were calculated for continuous variables (e.g. age); frequencies and proportions of categorical data (e.g. serotypes) were calculated.

### Reliability, Validity, and Objectivity

All tests were performed according to recognized, accredited standard operating procedures as well as to the instructions of the manufactures in the case where commercially available kits were used. Molecular size markers were used during agarose gel electrophoresis.

## RESULTS

### Study population

There were 50 *Cryptococci* isolates collected from different clinical specimens sent to the DGMT-Laboratory during the study period, June to October 2014 (5 months). Eleven (22%) isolates were from blood specimens and 39 (78%) from Cerebral Spinal Fluid (CSF).

The ages of the 41 patients analyzed ranged from 15 to 86 with the majority being between 35 and 45.

### Biochemical test for species

Urease slope was done to all 50 isolates. After a period of 24 hours incubation at 30°C, the color change was observed. The change of colorless broth to pink broth medium was confirming the presence of *C. neoformans* species **(figure 2)**.

**Figure 2:**
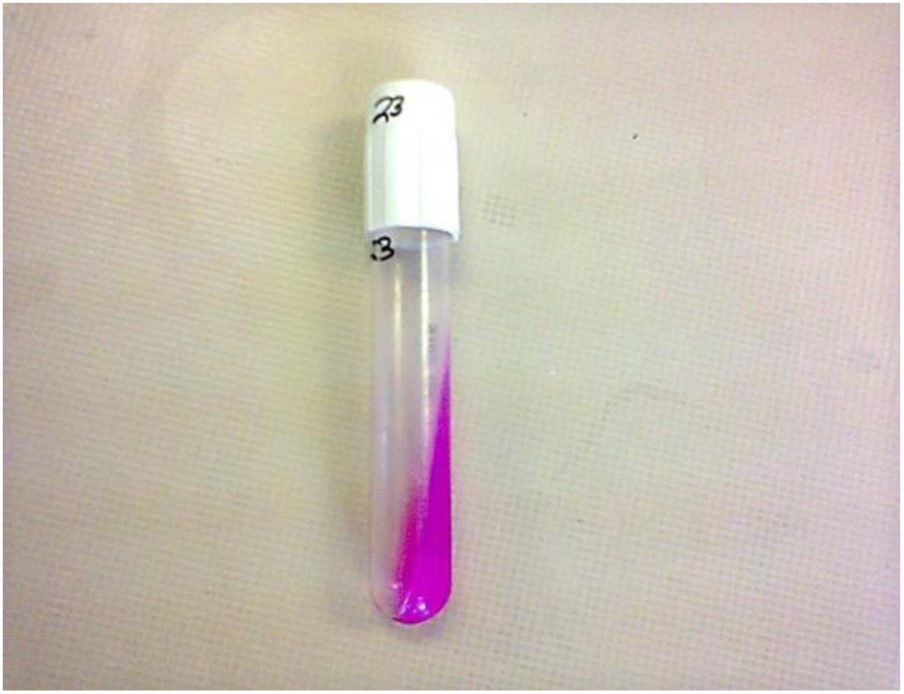
Urease slope of one of the isolates showing a colour change after 24 hours incubation.

### Susceptibility testing

After the incubation period of the three antifungals E-test strips (bioMérieux S.A., Marcy l’Etoile, France) which were fluconazole, voriconazole, and amphotericin-B, results were then read following the CLSI ^[24]^. The Minimum inhibitory concentration (MIC) values were read at the point of intersection between the zones of growth and the edge of the strip. The amphotericin-B was read at the point of complete inhibition (100%) as shown in **figure 3**, both fluconazole and voriconazole MICs were read at a point of significant inhibition of growth, about 80% reduction of growth as shown in **figure 4** and **5a-b**. MIC values were documented on a data collection sheet. The MIC values for fluconazole and voriconazole were interpreted in accordance with CLSI updated M27 breakpoints (2013) guideline and for amphotericin B, according to NCCLS M27 guideline^[24]^. These were interpreted as susceptible, intermediate and resistant.

**Figure 3:**
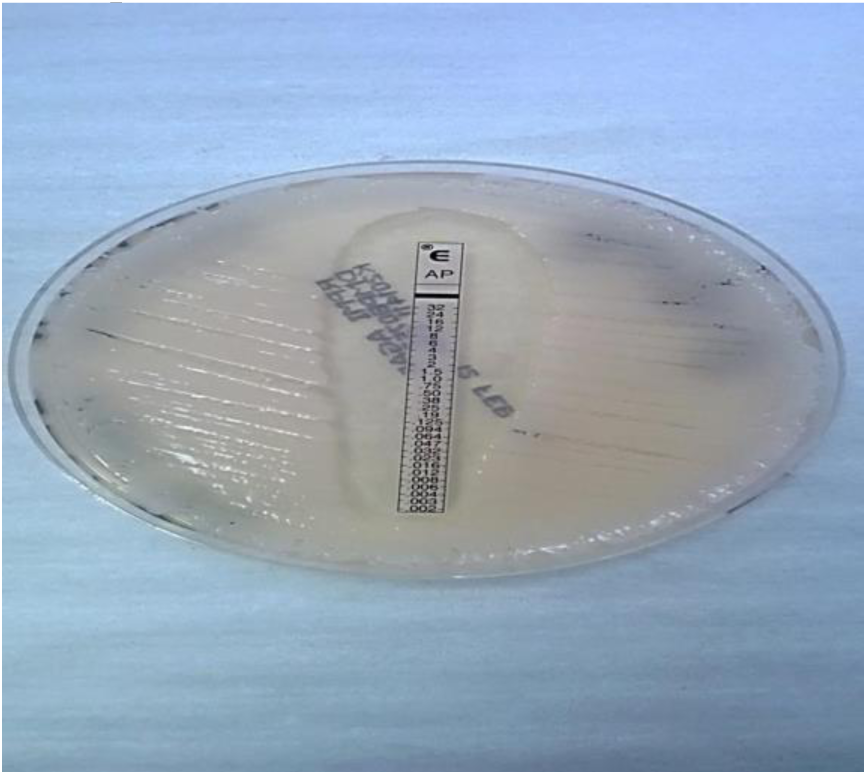
Amphotericin-B point of 100% inhibition of growth.

**Figure 4:**
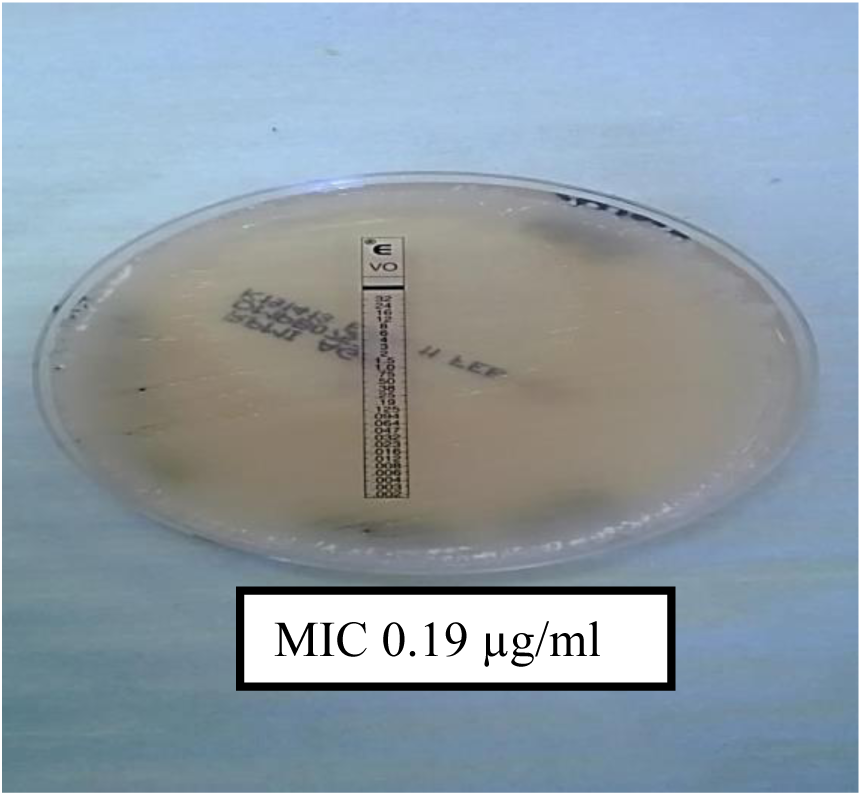
Voriconazole point of 80% inhibition of growth.

**Figure 5a:**
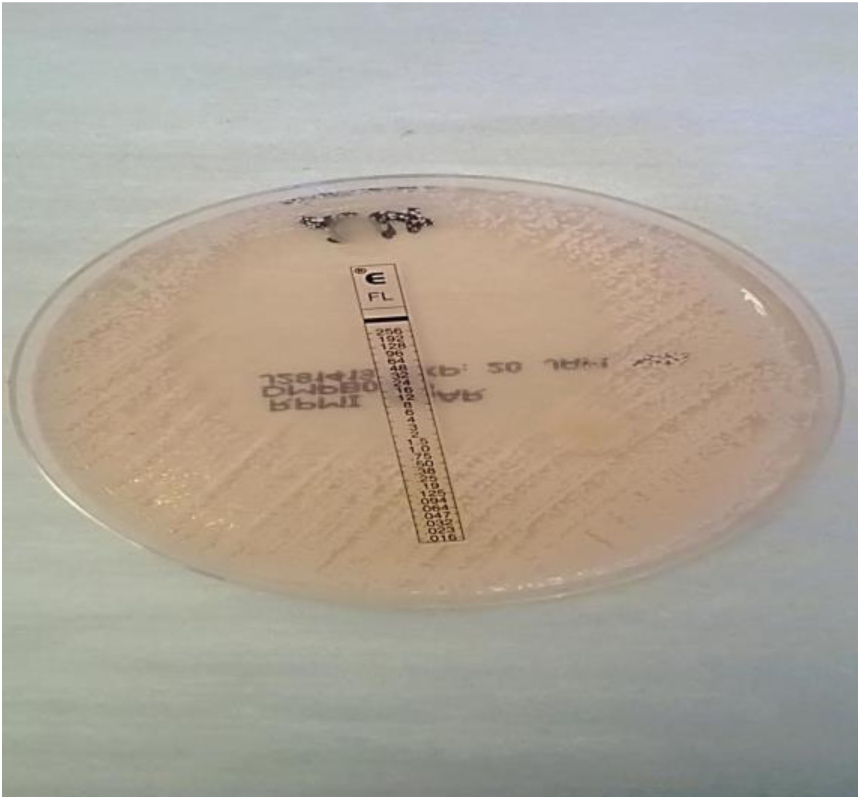
Fluconazole point of 0% inhibition of growth.

**Figure 5b:**
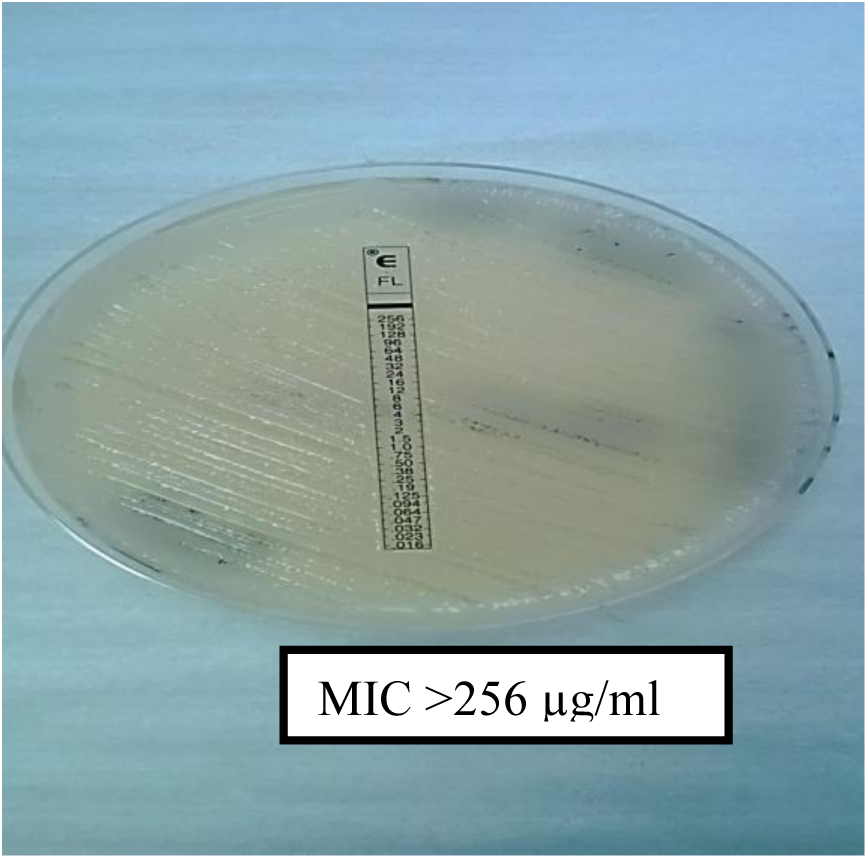
Fluconazole point of 80% inhibition of growth.

The MICs were determined in all isolates. Voriconazole and amphotericin-B were susceptible to all isolates as presented in **table 4** above.

**Table 4:** MIC’s of the isolates against common antifungals (N= 50)

| Antifungal drugs | Interpretation | MIC scales | Isolates numbers |
| --- | --- | --- | --- |
| **Fluconazole | Susceptible | ≤ 2 µg/mL | 13 (26%) |
|  | Intermediate | 4 µg/mL | 12 (24%) |
|  | Resistant | ≥ 8 µg/mL | 25 (50%) |
| **Voriconazole | Susceptible | $\leq 0.12 \mu\text{g/mL}$ | 50 (100%) |
| | Intermediate | $0.25 \mu\text{g/mL} - 0.5 \mu\text{g/mL}$ | 0 |
| | Resistant | $\geq 1 \mu\text{g/mL}$ | 0 |
| *Amphotericin-B | Susceptible | $\leq 0.5 \mu\text{g/mL}$ | 50 (100%) |
|  | Intermediate | - | 0 |
| | Resistant | $\geq 2 \mu\text{g/mL}$ | 0 |

\*interpretation according to NCCLS M27-A document 2000
\*\*interpretation according to CLSI M27-A document 2013

The agarose gel picture below, show representatives of PCR results on an agarose electrophoresis

**Figure:**
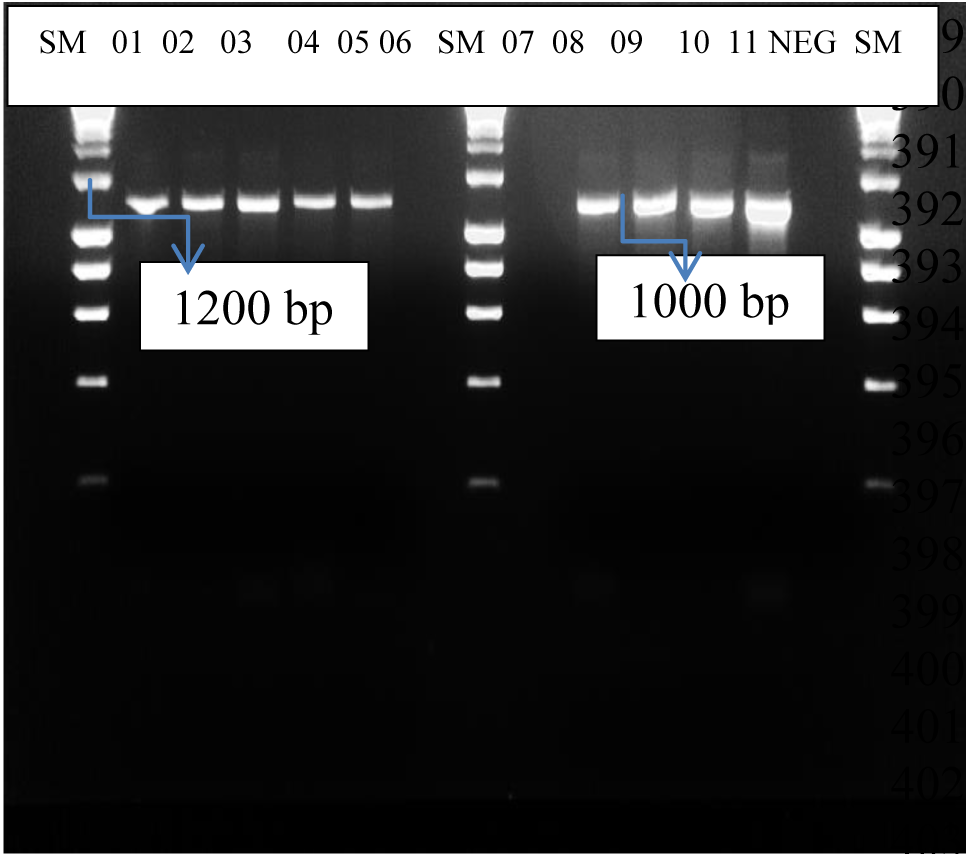
Representative Agarose gel electrophoresis. Where: Lane 1-11: Clinical isolates; Neg: Negative control; SM: 1000 bp (1 kd) (size markers); Lane number: 2-5; 8-11 represent serotype Aa mating genes; Lane number: 6-7 negative results.

## DISCUSSION

Resistance to antifungal agents used against cryptococcosis is globally reported^[9,16,29-31]^. In Africa, cryptococcosis epidemiology data is scarce but accumulated evidence in South Africa, makes it apparent that resistance development to commonly used antifungal agents is of concern^[8-10,25]^. Therefore, monitoring the susceptibility of these commonly used antifungal agents in different geographical areas is essential.

Data on circulating serotypes responsible for cryptococcosis in communities served by DGMT-Laboratory is not available. This study serves to profile the susceptibility and to identify the circulating serotypes of *Cryptococcus* at DGMT-hospital, in South Africa.

Based on our study, the susceptibility of the amphotericin-B, fluconazole, and voriconazole was profiled; resistance to fluconazole was of foremost concern **(table 4)**.

It was not surprising to see that half of our isolates were completely resistant to fluconazole. Our results were in keeping with multiple studies of diverse geographic areas, such that Arsenijevic et al (2014) in Serbia revealed 60% resistance of clinical isolates^[32]^, and that of 63% by Favalessa et al (2014) in West Brazil patients^[33]^. Furthermore, a South African report of Govender et al (2011) and (2013) showed 58% resistance to fluconazole^[9,25]^.

Fluconazole resistance is based on the *C. neoformans* mechanisms of action^[8,16,-18]^. The other factors that contribute to the recurrence of cryptococcosis among South African patients are limited access to treatment and inadequate treatment^[8-10,25]^.

Furthermore, isolates of our study were highly susceptible to voriconazole and amphotericin-B. Our findings were not different but comparable to the studies of Arsenijevic *et al* (2014)^[32]^, Govender *et al* (2011) and (2013), they all reported 100% susceptibility on voriconazole and amphotericin-B^[9,25]^. There was no cross-resistance between amphotericin-B, voriconazole, and fluconazole on in-vitro testing. It will, however, be important to assess this based on clinical outcome in patients.

Unfortunately, Amphotericin-B had no breaking-points according to CLSI updated M27 break-points document of 2013, we, therefore, interpreted our results in accordance with NCCLS M27-A guideline document (NCCLS M27-A guideline, 2000)^[24]^. Fortunately Govender ***et al*** (2011) also, however, indicated the challenges of performing susceptibility testing for amphotericin-B because of the absence of CLSI break-points^[9]^.

Molecular-based, our study confirmed that ***C. neoformans*** serotype A is predominant in our setting. Accumulated evidence showed that serotype A has been reported as more virulent and prevalent than the other serotypes^[32-36]^. Likewise, Lugarini et al (2008) in Brazil, reported a prevalence of 53% serotype A α-mating gene types circulating across the country^[34]^. A similar study by Favalessa et al (2014) in Midwest Brazil also reported serotype A making 63% of the isolates from HIV/AIDS patients^[33]^. Khayhan et al (2013) also confirmed serotype A as the most prevalent serotype in Asia Phayoa^[35]^. In our study, we didn’t manage to find the HIV status of our patients. Our study was in keeping with a systemic review study of Litvintseva et al (2011) which was conducted in African countries, reported serotype A specifically the α-mating gene types to account for 79% of the isolates^[36]^, and according to our study in South Africa, serotype A is the commonest circulating serotype across our setting, counting for 96% α-mating gene types.

Furthermore, our study showed that only a few isolates were confirmed to be serotype D α-mating genes type. Those few isolates were from patients over the age of 65. Duke University in Durham previously reported that serotype D is very rare and less information is documented about the distribution of this serotype^[37]^, whereas Feretzaki et al (2014) in India reported that serotype D requires very high inoculum to disseminate and cause infections like meningitis^[38]^. There is no information or data documented about the distribution of serotype D α-mating gene-types in South Africa and in our setting. Our two patients could have been more immune-compromised than the others because of their age. Furthermore, no study has been conducted according to our knowledge on serotypes and mating-genes in South Africa, Pretoria, DGMT-Hospital. Our results highlight the importance of properly treating cryptococcosis.

## CONCLUSION

*C. neoformans* serotype A is a predominant serotype in the area served by DGMT Laboratory, accounting for 96% of the isolates. Fifty percent of the isolates were resistant to fluconazole while 100% of those tested were susceptible to voriconazole and amphotericin-B, suggesting a lack of cross-resistance on in-vitro testing.

The study had several limitations such as low population number and financial constraints. However, because of high fluconazole resistance suggested, the study recommends the routine performance of susceptibility testing to fluconazole. Cross-resistance with voriconazole and amphotericin-B is to be evaluated further.

## Acknowledgments

We thank NHLS for assisting with isolates collections, and VLIR for reagents.

## Conflict of Interest

The authors declare no conflicts of interest with respect to authorship and/or publication of this article.

## Author Contributions

Conceived and designed the experiments: EZ Jiyane. Performed the experiments: EZ Jiyane, Analysed the data: EZ Jiyane, Contributed reagents/materials/analysis tools: VLIR, EZ Jiyane; Contributed to the writing of the manuscript: EZ Jiyane; critically reviewed the manuscript: EZ Jiyane, L Nemarude, Prof M Nchabeleng.

## Funding sources

All funds were received from VLIR.

